# Engineered biological neural networks on high density CMOS microelectrode arrays

**DOI:** 10.1101/2021.12.09.471944

**Authors:** Jens Duru, Joël Küchler, Stephan J. Ihle, Csaba Forró, Aeneas Bernardi, Sophie Girardin, Julian Hengsteler, Stephen Wheeler, János Vörös, Tobias Ruff

**Affiliations:** Laboratory of Biosensors and Bioelectronics, Institute for Biomedical Engineering, ETH Zürich, Zürich, Switzerland; Stanford University, Cui Laboratory, 290 Jane Stanford Way, Stanford, CA, USA

**Keywords:** Bottom up neuroscience, in vitro, CMOS, microelectrode arrays, engineered networks, PDMS microstructures

## Abstract

In bottom-up neuroscience, questions on neural information processing are addressed by engineering small but reproducible biological neural networks of defined network topology *in vitro*. The network topology can be controlled by culturing neurons within polydimethylsiloxane (PDMS) microstructures that are combined with microelectrode arrays (MEAs) for electric access to the network. However, currently used glass MEAs are limited to 256 electrodes and pose a limitation to the spatial resolution as well as the design of more complex microstructures. The use of high density complementary metal-oxide-semiconductor (CMOS) MEAs greatly increases the spatiotemporal resolution, enabling sub-cellular readout and stimulation of neurons in defined neural networks. Unfortunately, the non-planar surface of CMOS MEAs complicates the attachment of PDMS microstructures. To overcome the problem of axons escaping the microstructures through the ridges of the CMOS MEA, we stamp-transferred a thin film of hexane-diluted PDMS onto the array such that the PDMS filled the ridges at the contact surface of the microstructures without clogging the axon guidance channels. Moreover, we provide an impedance-based method to visualize the exact location of the microstructures on the MEA and show that our method can confine axonal growth within the PDMS microstructures. Finally, the high spatiotemporal resolution of the CMOS MEA enabled us to show that we can guide action potentials using the unidirectional topology of our circular multi-node microstructure.

## INTRODUCTION

How networks of biological neurons process, store and retrieve information remains an unanswered question to this day, despite a vast amount of tools available to establish neural interfaces. Patch-clamp technique has enabled highly sensitive recordings and stimulation of isolated neurons down to the level of single ion channels (Johansson and Arhem, 1994). However, the patch-clamp technique requires a high level of technical expertise and the requirement for sequential patching of each individual neuron highly restrict the throughput and application of the technique.

On the other hand, functional magnetic resonance imaging (fMRI) allows for an analysis of neuronal dynamics in large areas of brain tissue at the cost of limited temporal and spatial resolution (Lewis et al., 2016). To overcome those limitations, microelectrode arrays (MEAs) have been developed and applied to study the behavior of neural networks *in vitro* (Bakkum et al., 2008), *ex vivo* (Ha et al., 2020), and even *in vivo* (Wei et al., 2015). Such devices allow for simultaneous extracellular recording and stimulation at up to several thousand electrodes across the spatial extent of the electrode array. However, in all cases the vast amount of interference of the measured neural circuit with other neurons and the high degree of variability between brains and brain explants of even the same species prevented true understanding on how even small neural networks operate. A true understanding of neural networks would mean that we can combine neural elements with defined connectivity to generate a network with a predictable output. This approach to reverse engineer the brain from its fundamental unit - the single neuron - is termed bottom-up neuroscience. To enable bottom-up neuroscience research, it is essential to gain electrical modulatory access to individual neurons and have full control over the network topology. A distinct network topology can be achieved by confining the adhesion sites of neurons and influence the growth-direction of axons by chemical surface patterning (Jang and Nam, 2012), the usage of microstructures (Renault et al., 2016; Ming et al., 2021), and other techniques involving electrokinetic (Honegger et al., 2013) or optical guidance (Stevenson et al., 2006). From all mentioned techniques, the physical confinement of cells in microstructures is considered the most reliable method to establish engineered biological neural networks *in vitro* (Aebersold et al., 2016).

Using polydimethylsiloxane (PDMS) microstructures, any network topology consisting of multiple compartments interconnected with thin axon guidance channels can be generated. Thus, PDMS microstructures allow full control over topology-relevant parameters such as the distance and directionality of connections between neurons. Previously, this method was used to implement engineered biological neural networks on 60-channel glass MEAs. The high compatibility of PDMS microstructures with the flat surface of standard glass MEAs allows the alignment of channels and microcompartments on top of each of the 60 electrodes (Forró et al., 2018). The combination of MEAs with PDMS microstructures defines the connectivity of small neural populations and allows for selective electrical access to the neural culture. One key limitation of this approach is the limited number of electrodes that restricts PDMS design options and requires a tedious process of structure-to-electrode alignment. More importantly, the high electrode pitch in the order of 100 µm present in these MEAs only allows the detection of electrical activity of multiple axons growing within the channels of the microstructure (Figure 1A).

**Figure 1.**
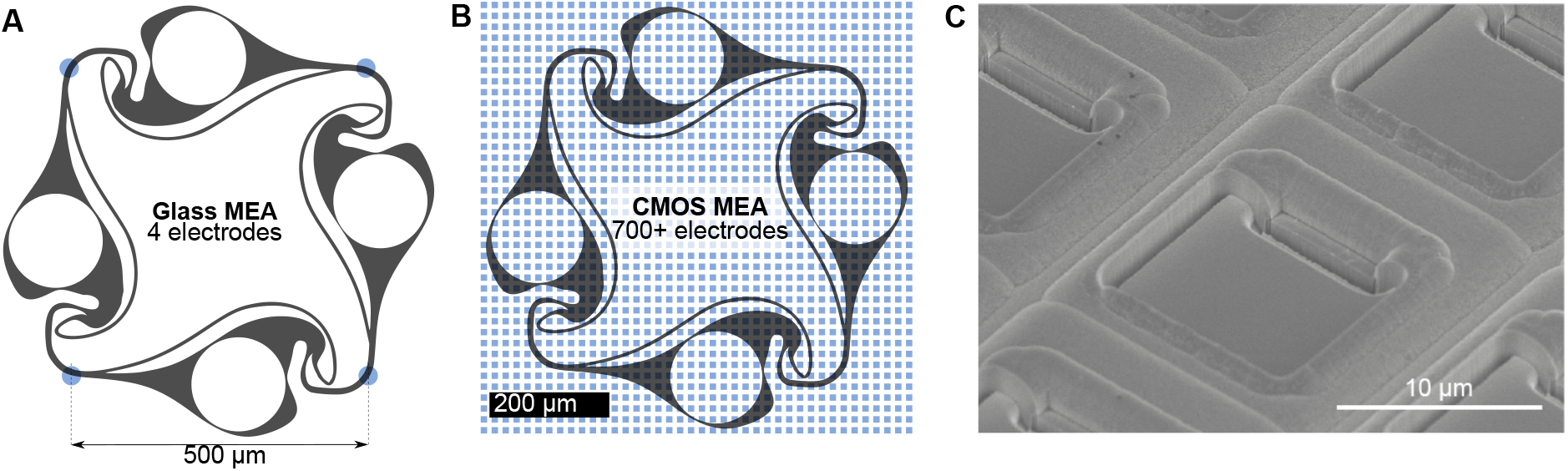
Comparison of recording electrode densities on 60-channel glass MEAs and CMOS MEAs. **(A)** Commonly used 60-channel glass MEAs allow for simultaneous recording from an engineered biological neural network at 4 electrodes (blue dots) located at the microstructure channels. **(B)** The same microstructure on a CMOS MEA is covered with more than 700 electrodes, allowing for both axonal and somatic signal acquisition. **(C)** Electron micrograph of the chip surface. While the glass MEAs are flat, the surface of the CMOS MEA is non-planar due to recessed electrodes and trenches between columns of electrodes. Such a surface topology impairs the adhesion of the PDMS microstructure to the surface compromising its cell and axon guidance characteristics.

In recent years, the advent of complementary metal-oxide-semiconductor (CMOS) microelectrode array technology has led to a massive expansion in the number of electrodes per array (Obien et al., 2015). By including filtering and amplification stages directly on the chip in the vicinity of each microelectrode, wiring of the electrode to external signal processing elements becomes less susceptible to noise and high electrode densities can be realized (Heer et al., 2006). Commercially available chips offer up to 26400 electrodes confined within a sensing area of 3.85 × 2.10 mm^2^ (Müller et al., 2015). The high electrode density enables action potential propagation tracking with subcellular resolution (Bakkum et al., 2013) as well as the stimulation of single neurons (Ronchi et al., 2019). Upcoming, experimental stage CMOS MEAs offer the opportunity to record the electrical activity from neural cultures at even up to 19584 recording sites simultaneously (Yuan et al., 2021).

Combining CMOS arrays with PDMS microstructures would enhance the number of recording sites by orders of magnitude (Figure 1B) with respect to 60-channel glass MEAs and enable high resolution recording from neurons of defined connectivity. In the past, CMOS MEAs were used in combination with PDMS microstructures to isolate cell bodies from axons and record axonal signals along microchannels running in parallel to electrode columns of the CMOS MEA (Lewandowska et al., 2015). However, the non-flat surface of the CMOS chips (Figure 1C) has so far prevented the use of more complex PDMS microstructures together with CMOS arrays, due to axons escaping the PDMS confinement through the ridges between the columns of microelectrodes. Here, we present a method to glue PDMS microstructures with feature sizes down to 5 × 4 µm^2^ onto the uneven surface of a non-planar CMOS MEA and show that this method confines cell bodies and axons within the microstructures. Moreover, we developed an impedance-based protocol to identify the exact location of all microstructure elements, which enables selective routing of almost all underlying electrodes. Finally, we show that our method enables us to track the spatiotemporal spread of action potentials within our directional circular four-node microstructure networks at a very high signal-to-noise ratio.

## MATERIALS AND METHODS

### Microelectrode Array

A CMOS technology based high-density microelectrode array (MaxOne) from Maxwell Biosystems (Zürich, Switzerland) was used for all experiments. The MEA provides 26400 electrodes in a 3.85 × 2.10 mm^2^ large sensing area with an electrode pitch of 17.5 µm (Müller et al., 2015). An almost arbitrary combination of up to 1024 electrodes can be recorded from simultaneously. Electrode patches of 23 × 23 electrodes can almost always be routed completely and used as a dense recording site. In case larger electrode patches are selected, the number of routed electrodes is maximized using a routing algorithm that aims to maximize the number of electrodes that are connected via the switch-matrix to the 1024 available amplifiers (Frey et al., 2010). Data is acquired with a sampling rate of 20 kHz (50 µs temporal resolution).

### PDMS microstructures

PDMS microstructures for cell and axon guidance were designed in a CAD software and fabricated through a soft lithography process by Wunderlichips (Zürich, Switzerland). The microstructure consists of 15 substructures, i.e. 15 individual networks. The layout of such a network is shown in Figure 1A and B. Each network consists of 4 nodes, in which the neurons can enter through a 170 µm large hole during the seeding process. The nodes are connected with microchannels of size 10 × 4 µm that are impenetrable for the cell body but accessible by axons and dendrites. The shape of the structure was designed to promote directional growth of axons in a clockwise fashion. Microchannels of size 5 × 4 µm running in parallel to the nodes redirect axons growing in anti-clockwise direction. The design principle of the single stomach shaped nodes is explained in detail in a previous publication (Forró et al., 2018).

### Microelectrode array preparation and PDMS microstructure adhesion

Both new MEAs as well as MEAs that have been previously used to record from random (i.e. without PDMS microstructures) cultures were used. In order to promote cell adhesion, 100 µL of 0.1 mg/mL Poly-D-Lysine (PDL, Sigma-Aldrich, P6407) was pipetted onto the electrode area of the MEA and incubated at room temperature for 30 min. Unbound PDL was rinsed away in three washing steps with deionized water. After blow-drying the MEA surface with N2, the PDMS microstructure was transferred onto the MEA with an intermediate gluing step (Figure 2). For this, PDMS with the standard 1:10 weight ratio of crosslinker to base (SYLGARD 184 Silicone Elastomer Kit, Dow Chemical) was diluted with hexane (Sigma-Aldrich, 34859) in a volume ratio of 1:9. The mixture was transferred to a silicon wafer and spin-coated at 2000 rpm for 60 s at room temperature. A PDMS microstructure was cut and placed carefully on the spin-coated PDMS thin film with tweezers in order to coat the bottom of the PDMS microstructure with a thin layer of uncured PDMS. The microstructure was then placed onto the center of the MEA and cured for 2 h at 80°C. To fill the hydrophobic microchannels, 1 mL of phosphate buffered saline (PBS, ThermoFisher, 10010023) was pipetted onto the MEA. Then, the MEA was desiccated for 10 minutes. Emerging bubbles stuck to the PDMS surface were removed by inducing a flux with a pipette. This desiccation process was repeated three times.

**Figure 2.**
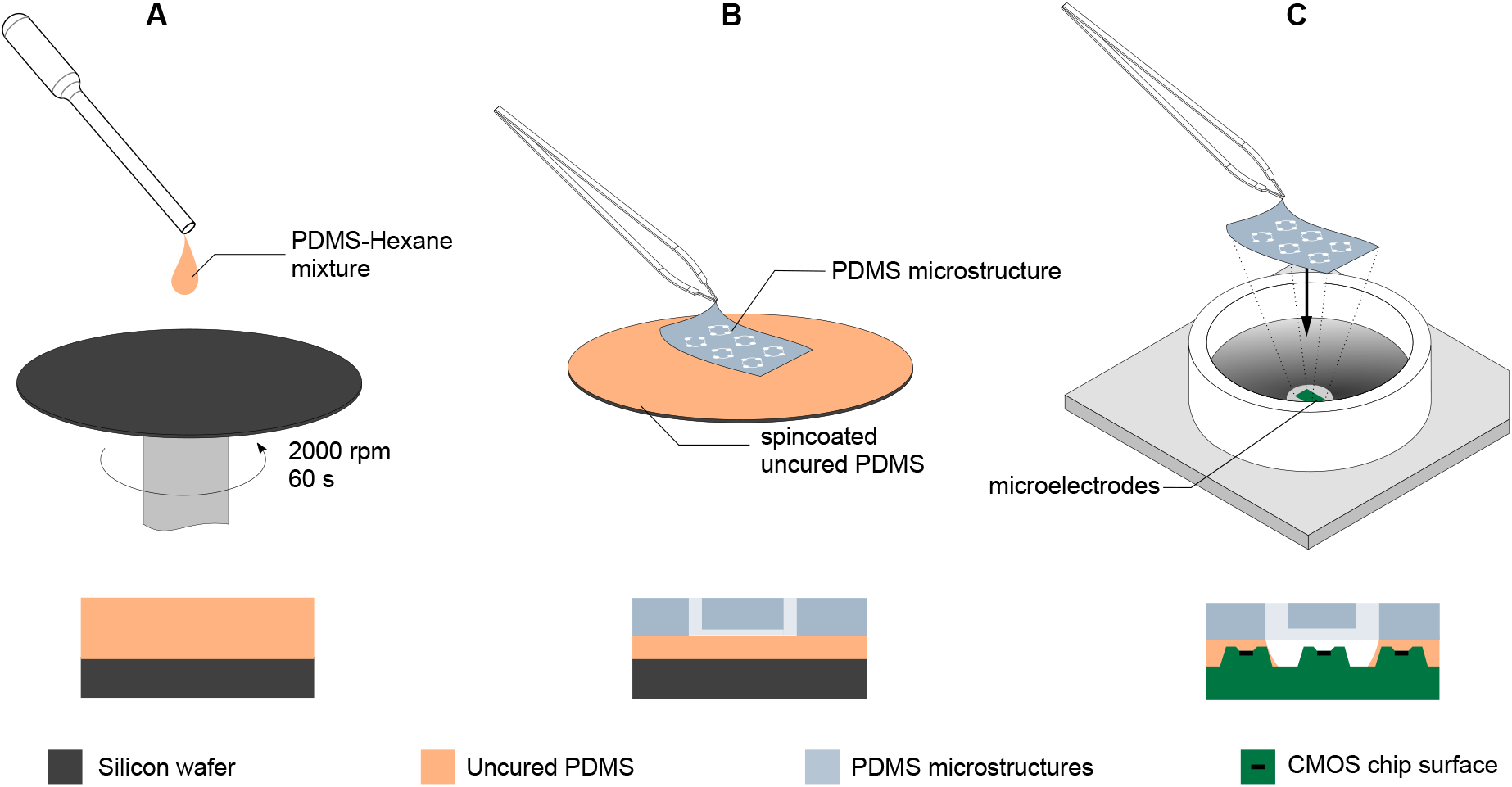
Transferring PDMS microstructures to a CMOS MEA using an intermediate gluing step with uncured PDMS. **(A)** Uncured PDMS is diluted with hexane in a 1:9 volume ratio and spin coated on a wafer at 2000 rpm for 60 s. **(B)** The PDMS microstructure is placed on the spin coated PDMS thin film without releasing it and lifted after contact was established, leaving a thin layer of uncured PDMS on the bottom side of the PDMS microstructure. **(C)** The PDMS microstructure with the intermediate layer of uncured PDMS is placed on the CMOS MEA and subsequently moved to an oven at 80 °C for 2 h to allow the intermediate PDMS layer to fully cure and create a tight seal between the surface of the MEA and the PDMS microstructure.

### Locating the PDMS microstructure on the MEA

To determine the location of the electrodes underlying the PDMS microstructure, an impedance map was generated using a custom Python script. A sinusoidal electrical signal (16 mV peak-to-peak voltage, 1 kHz frequency) was generated using an on-chip function generator. The sinusoidal signal was applied to the circumferential reference electrode of the MEA and the received signal at the microelectrodes was recorded (Figure 3A). Since the MEA provides only 1024 readout channels, the signals were recorded using a total of 25 contiguous routing patterns. Electrodes with an x-coordinate > 210 were discarded in the analysis due to access limitations of the switch matrix that result in complicated routing patterns that need to be generated. A recording was done on each pattern for 5 s and the peak-to-peak amplitude of the received signal on each microelectrode was determined. Electrodes insulated by the PDMS microstructure showed highly attenuated signals, while electrodes outside of or within the openings of the microstructure showed signals with only minimal signal attenuation. By plotting the peak-to-peak amplitudes in a 120 × 220 element matrix, an impedance map can be generated (Figure 3B) and saved for further usage in custom recording software to determine the location of electrodes covering individual networks within the PDMS microstructure.

**Figure 3.**
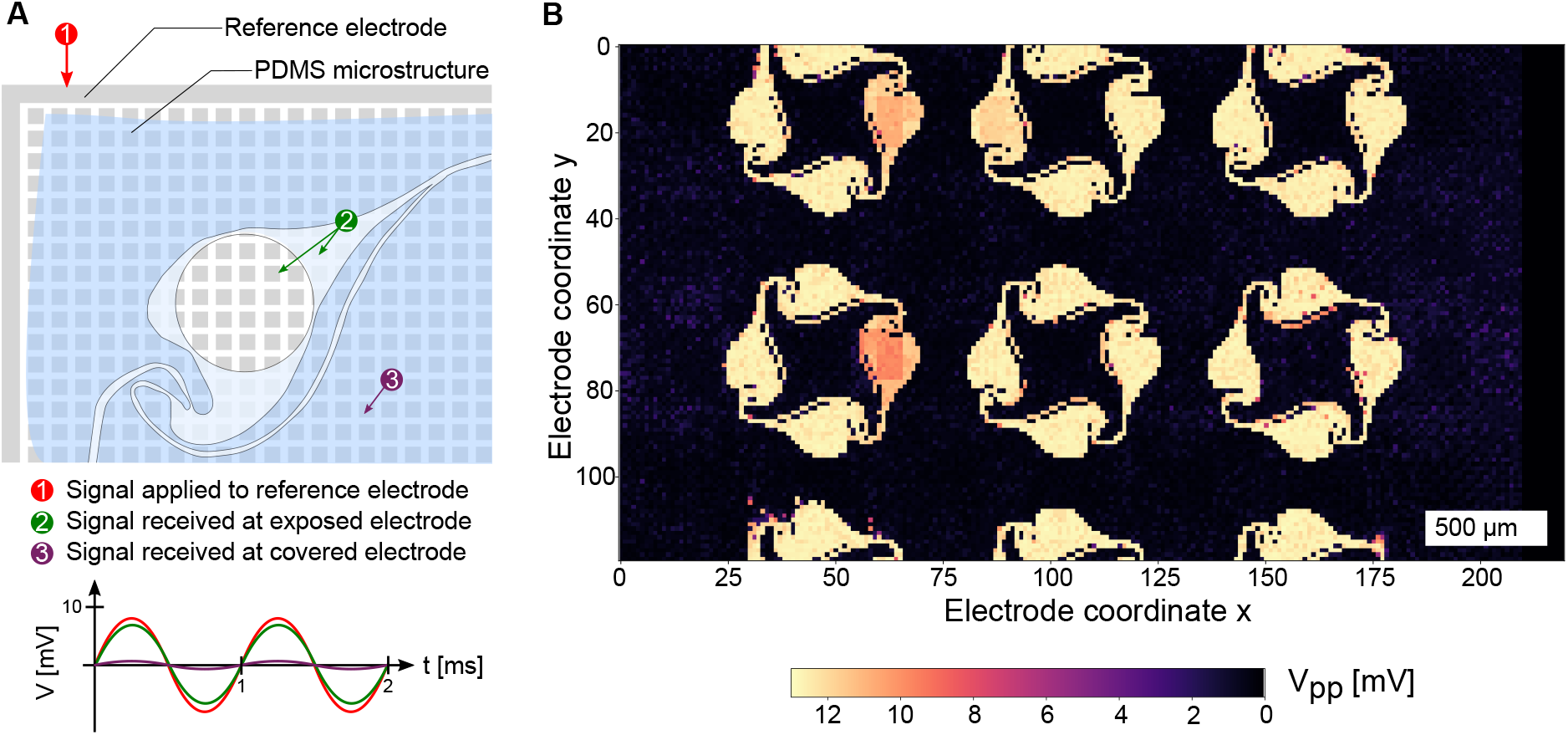
Detection of the PDMS microstructure on the CMOS MEA. **(A)** Applying a sinusoidal signal on the reference electrode with an on-chip signal generator yields an undamped signal on exposed electrodes and a highly attenuated signal on covered electrodes. **(B)** Repeating the procedure on 25 routing patches across the MEA reveals the location and openings of the PDMS microstructure.

### Cell seeding

The PBS used in the desiccation and microstructure location step was aspirated and replaced by Neurobasal medium (21103049) with 2 % B27 supplement (17504001), 1% GlutaMAX (35050-061) and 1% Pen-Strep (15070-063, all from ThermoFisher). The medium was warmed up and passively pH adjusted in the incubator for at least 30 min. Then primary cells from the cortex or hippocampus of E18 Sprague-Dawley rat embryos (Janvier Labs) were seeded onto the CMOS MEA with seeding densities ranging from 125,000 to 250,000 cells/MEA. The following dissociation protocol was adapted from Forró et al. (2018). The cortices were dissociated for 15 min using a filtered and sterilized solution of 0.5 mg/mL papain (Sigma-Aldrich, P4762) with 0.01 mg/mL deoxyribonuclease (Sigma-Aldrich, D5025) in PBS, supplemented with 0.5 mg/mL bovine serum albumin (Sigma-Aldrich, A7906) and 10 mM D-(+)glucose (Sigma-Aldrich, G5400). The supernatant was removed and replaced with Neurobasal medium three times before a gentle trituration step. The suspended cells were then pipetted onto the MEA and allowed to settle onto the surface for 30 min in the incubator. Afterwards, a pipette was used to recirculate the cells that did not fall into the microstructure opening, by inducing a flux within the culture medium on the MEA. The resuspended cells were given another 30 min to settle. To remove cells that landed on top of or outside the PDMS microstructure, the culture medium was aspirated away and 1 mL of Neurobasal medium was added to the MEA. Half of the medium was replaced on day *in vitro* (DIV) 1 and twice a week from then on. In order to slow down medium evaporation in the incubator, a custom-made polytetrafluoroethylene (PTFE) ring covered by a fluorinated ethylene-propylene membrane (ALA MEA-MEM-SHEET, ALA Scientific Instruments Inc.) was designed to be placed on top of the culture chamber of the MEA.

### Network selection and recording

The impedance map can be read into a home-built Python program with a user-friendly graphical interface. The location of electrodes covering individual networks within the microstructure can be determined by thresholding a region of the impedance map. The information is then sent to the FPGA, which runs a proprietary algorithm to maximize the number of routed electrodes. The FPGA is connected to the headstage in which the MEA can be inserted. The spontaneous activity of the engineered neural network can be recorded for a user-specified amount of time and the data is stored in an hdf5 file on the connected computer.

### Raw data processing

The raw data containing hdf5 file was processed using a home-built Python processing pipeline. First, the data was filtered using a 2nd order butterworth filter (high pass) with a cutoff frequency of 200 Hz. A peak-detection algorithm determined the location of action potentials by determining the times at which the signals on each individual electrode surpassed 5 times the standard deviation σ of the filtered signal. In order to prevent that the negative and positive phase of the spike were counted as two individual spikes, a window of 100 samples (5 ms) was selected in which local maxima in the signal were discarded. Spike average maps were generated by creating the average spike shape at each electrode. For this, the filtered signal was cropped 75 frames (3.75 ms) before and after each detected spike and the cropped signal was subsequently stored and averaged. The average spike shape was then plotted at the location of the corresponding electrode. Signal propagation movies were generated by plotting the signal at each time step within a specified time window. The plots were then combined to yield a video to show the propagation of signals within the network.

### Cell culture and microstructure imaging

The viability of the cell culture and the guidance properties of the PDMS microstructure were investigated using a confocal laser scanning microscope (Olympus FluoView 3000). To get a fluorescent staining of the live and dead neurons, the cells were incubated for 30 min with CellTracker Green CMFDA (ThermoFisher, C7025) and ethidium homodimer (ThermoFisher, E1169) both at a concentration of 0.5 µM. The culture medium was washed away and replaced with fresh medium. To image the culture in an inverted microscope, a coverslip was placed in the culture medium on top of the MEA sensing area and the medium was aspirated around the coverslip, leaving behind a thin film of medium secured by a coverslip when flipping the MEA. The MEA was inserted into the headstage, which was placed in a custom home-made aluminum holder and imaged using a 10x objective. The procedure is illustrated in the supplementary Figure S1. In order to assess the adhesion of the PDMS microstructure to the MEA surface, the microstructure was removed from the MEA and cleaned from cell debris using tergazyme. Afterwards, it was placed on a coverslip and imaged with phase contrast imaging.

### Electron microscopy of the chip surface

Electron microscope images were taken with a scanning electron microscope (SEM, Magellan 400, FEI Company, USA). Prior to imaging, the sample was sputter-coated with a thin Pt-Pd layer (3 nm) using a vacuum coating system (CCU-010, safematic GmbH, Switzerland). All SEM images were taken with an acceleration voltage of 5 kV at a 45 degree tilt angle.

## RESULTS

### PDMS thin film stamp transfer enables PDMS microstructure bonding to non-flat CMOS MEAs

Both electrical recordings as well as the fluorescent images of the cultures on the CMOS MEAs reveal that the intermediate PDMS gluing step is crucial for successful PDMS microstructure adhesion, network formation without channel clogging, and axonal growth within the microchannels. A too thin layer of the spincoated diluted PDMS film results in poor adhesion of the PDMS microstructure to the CMOS MEA surface and was observed when using diluted heptane instead of hexane. While neurons were able to enter the PDMS microstructure from above through its nodes during seeding, no axonal guidance of the PDMS microstructure was present and axons escaped following the non-planar topology of the chip. The brightfield image of the microstructure after removal from the MEA surface reveals that while all channels are open, only a slight imprint of the MEA surface is visible on the microstructure. This implies that only a small amount of the PDMS-heptane mixture entered the ridges of the CMOS MEA without sealing them from axon access (Figure 4A). The highest success rate was achieved when PDMS was mixed with hexane in a 1 to 9 ratio and no pressure was applied onto the microstructure after it was placed on the CMOS MEA. An exemplary fluorescent image reveals full connectivity in the microchannels interconnecting the nodes (Figure 4B). The brightfield image of the microstructure shows that the microchannels interconnecting the nodes are indeed open, while the extremely thin microchannels (5 µm) running in parallel to the nodes are clogged. When pressure was applied to the microstructure after it was transferred to the CMOS MEA, the thin PDMS film was able to fill the ridges of the CMOS array but unfortunately also clogged the microchannels of the PDMS microstructure. As a result, the neurons seeded into the microstructure were not able to extend axons into the microchannels of the PDMS microstructure. The exemplarly brightfield image confirms clogging of all microchannels as well as a strong imprint of the MEA surface onto the microstructure (Figure 4C). Clogging of the microstructure microchannels is also visible, when the thickness of the spincoated PDMS film on the wafer was increased at higher dilution ratios of PDMS-to-hexane.

**Figure 4.**
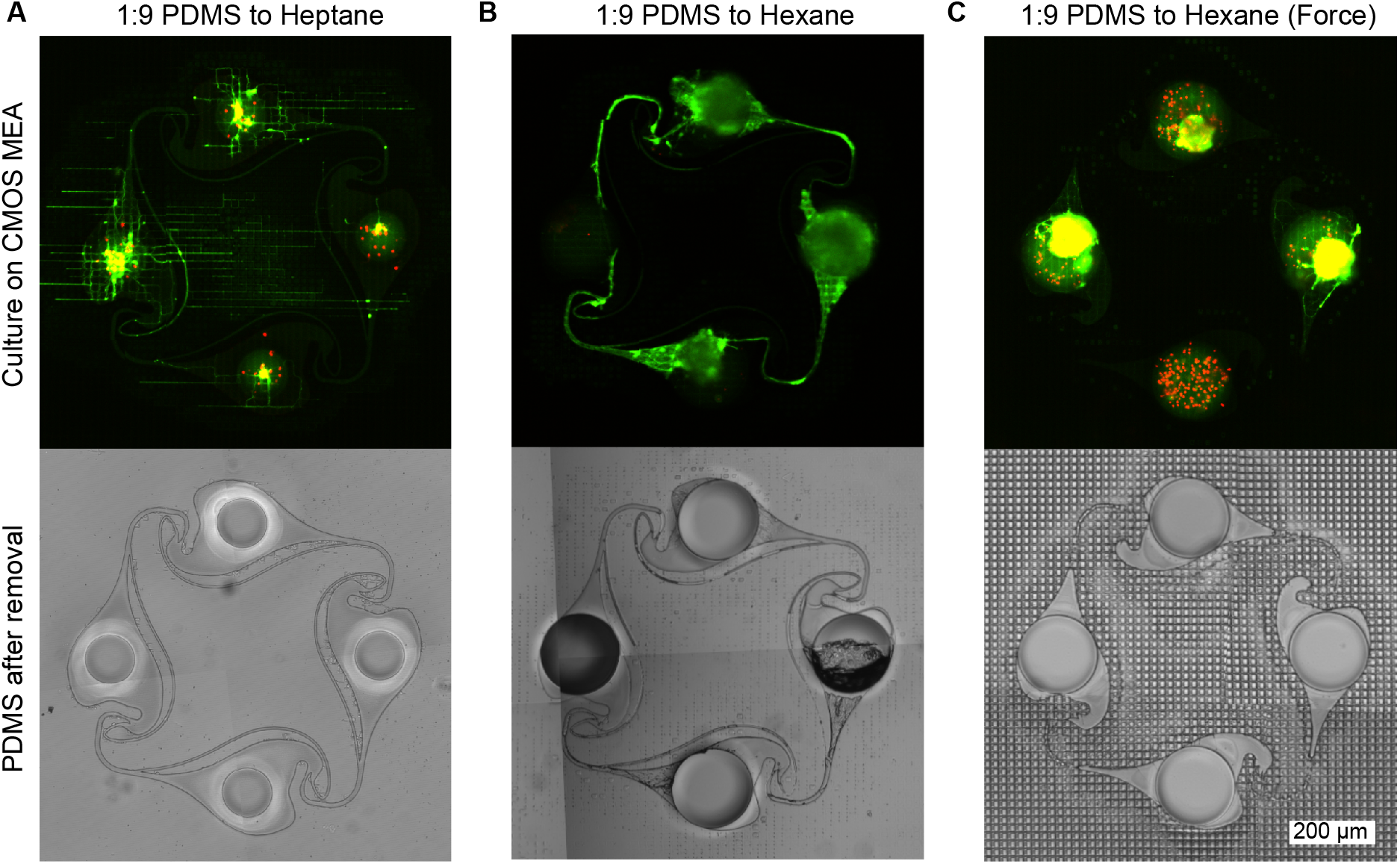
PDMS microstructure adhesion on the CMOS MEA surface depending on intermediate PDMS film quality. **(A)** The adhesion of the microstructure is low when heptane is used instead of hexane as a dilutant, since the same spincoating parameters resulted in a thinner PDMS film on the wafer. Consequently, less PDMS is transferred by the microstructure onto the CMOS array resulting in incomplete sealing of the trenches on the chip surface. Therefore, axons can escape the PDMS microstructure more easily and instead follow the topology of the MEA surface rather than the channels of the microstructure. After removing the microstructure from the MEA, a phase contrast image of the microstructure reveals that the MEA surface left almost no imprint on the PDMS microstructure, confirming that the intermediate PDMS layer did not fill the trenches of the MEA surface. **(B)** Using a spincoated 1:9 PDMS:Hexane film yields the best microstructure adhesion and axonal guidance. After removing the microstructure, the imprint of the surface of the CMOS MEA is visible. While some clogging is observable in the thin channels, the large channels interconnecting the nodes remain open. **(C)** When pressing the PDMS microstructure onto the CMOS MEA or using less diluted PDMS solutions, clogging is observed in all channels and only little axonal outgrowth is possible. The phase contrast image of the microstructure confirms the clogging, as the MEA surface imprint is also visible in the channel regions.

### Quality assessment of the PDMS microstructure bond by impedance map and neural activity measurements

In total, 16 PDMS microstructures were attached to different CMOS MEAs using the protocol mentioned in the methods section. Culture images, activity recordings, and impedance maps were obtained for 14 chips. The impedance maps obtained from these chips show a total of 47 individual networks, that were fully covered by microelectrodes, i.e. all four nodes and interconnecting microchannels of the PDMS microstructures were completely located on the sensor area. Out of these 47 covered networks, 11 networks showed the full shape of the used microstructure design and no sign of clogging of the 10 µm wide inter-node channels on the impedance map (23.4 %). A network is considered open if all electrodes positioned below an inter-node channel have at least one neighboring non-covered electrode. On 6 out of these 14 MEAs (42.9 %), neural activity was measured in at least one microchannel before or at DIV 14. In total, 9 out of the 47 independent networks (19.1 %) showed this behavior. Full connectivity - i.e. activity measured in all four microchannels of a four node structure without any clogged channel - was observed in 5 networks (10.6 %) on 2 different CMOS MEAs (14.3 %). All culture images and impedance maps are available in the supplementary information (Figures S2 and S3).

### Unidirectional action potential propagation in a circular four-node network

One of the fully interconnected networks is shown in Figure 5A. It is evident that neurons are present in all four nodes and axons were able to grow from one node into a neighboring one, creating a fully connected engineered biological neural network, which is entirely covered by microelectrodes. From the impedance map, it was determined that the whole network is covered by a total of 760 microelectrodes. All electrodes were selected for routing and 685 were successfully connected to the amplifying stage for signal acquisition (90.1 %). The recorded spikes are extremely large (Figure 5B), which is common for recordings from axons confined in microchannels (FitzGerald et al., 2008; Pan et al., 2014). Spikes are adequately detected using the spike detection algorithm described in the methods section. A spike frequency map confirms that the highest activity is detected in these microchannels, while signals within the node are less prominent in the measurement. In addition to determining spike locations and spike times, the spike shape can also be recorded. The average spike shape on every electrode was determined and plotted at the corresponding electrode location (Figure 5C). In general, the spike average and frequency map coincides well with the culture image shown in Figure 5A. The high temporal resolution of the CMOS MEA allows to track the propagation of action potential within a network. The chosen PDMS microstructure was designed to promote clockwise directionality of axonal growth and hence action potential propagation, which was also observable in this network (Figure 5D). A video demonstrating the propagation of action potentials within the network can be found in the supplementary information (Movie S1).

**Figure 5.**
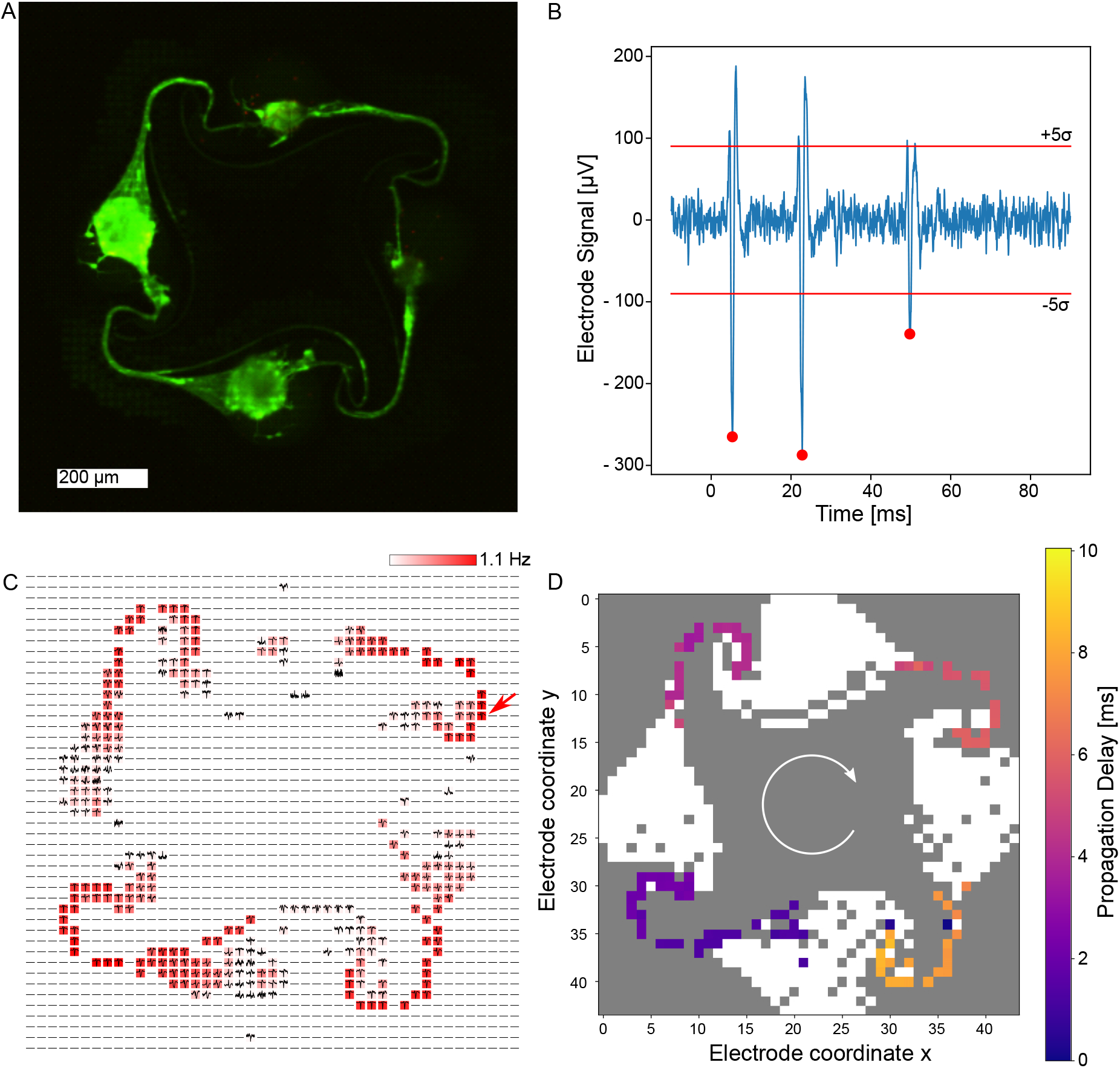
Electrical recordings of cortical neurons growing within a PDMS microstructure at DIV 14. **(A)** Fluorescently stained neurons imaged at DIV 31 reveal high viability, axonal growth within the microstructure channels and a full intra-node connectivity. **(B)** Recording of a single electrode shows a high signal-to-noise ratio. The 5σ threshold is indicated as well as spikes that were detected using the spike detection method described in the methods section. **(C)** Spike average and frequency map over 60 s on all routed electrodes within the network with color coded spiking frequency. It is evident that the spiking frequency is highest in the channels of the microstructure. The arrow indicates the electrode location at which the signal in **(B)** stems from. **(D)** A temporal analysis of spontaneous activity within a 10 ms time window indicates clockwise signal propagation.

## DISCUSSION

Combining PDMS microstructures with glass MEAs enables axon guidance across electrodes and the formation of any desired network topology. In order to increase the electrode density, we aimed to transfer this technology to high density CMOS microelectrode arrays. However, mounting PDMS microstructures onto CMOS MEAs has so far resulted in axons escaping from the channels due to the non-planar surface of the CMOS chip.

In this study, we demonstrated a stamp transfer method using hexane diluted PDMS (1:9, PDMS:Hexane) to bond PDMS microstructures onto CMOS chips. Successful stamp transfer depended on a thin PDMS film to be transferred from the PDMS-wafer contact surface onto the CMOS chip without clogging the PDMS microchannels, while still providing excess PDMS to fill the ridges on the CMOS to prevent axonal escape. Successful stamp transfer of PDMS microstructures with 10 µm tall channels onto flat surfaces has already been shown by Wu et al. (Wu et al., 2005). In their work, a 400 nm thick PDMS film was ideal for a permanent bond without clogging the channels. Similarly, we kept a constant spin-coating speed and time and regulated the PDMS film thickness by decreasing the viscosity of PDMS. Reducing the viscosity with hexane allowed us to significantly reduce the PDMS film thickness and is a common strategy to mold PDMS into submicrometer structures (Lee et al., 2019; Con and Cui, 2013).

Small axon guidance channels have the advantage that axons will not spontaneously turn around and grow in reverse direction. Moreover, they enable a wider repertoire of network architectures in a more compact arrangement. We have successfully bonded channels with sizes down to 10 × 4 µm onto the CMOS chip. However, channels smaller than 10 µm width frequently got clogged. The stamp transfer method relies on a balance between transferring a layer of PDMS that is thin enough to not clog the small PDMS microchannels but sufficiently thick to fill and seal the ridges on the CMOS array. Thus, we believe it would be difficult to further optimize our method for smaller channels. We believe that the width of the 5 × 4 µm channel should not have an effect on the redirection capabilities of axons and hence increasing the size of these thin channels in future designs may prevent clogging.

In our previous publication (Forró et al., 2018) we have already demonstrated how the assymetric design of the PDMS microstructures translates into a more directional signal propagation. The limited number of electrodes on the glass MEA restricted the measurement of spikes to few locations between the nodes resulting in a low electrode-to-neuron ratio. This makes spike sorting more difficult and assumptions on signal propagation speed and synaptic delays had to be made to assess how the signal travels through the microstructure (Forró et al., 2018). The high density of electrodes and the increased signal-to-noise ratio of the CMOS arrays however enables us to identify the origin and measure the propagation direction and speed of action potentials within each circuit (Emmenegger et al., 2019). Although we have not done any spike sorting in this work, the higher ratio of electrodes per neuron should facilitate classification of spikes to each neuron (Diggelmann et al., 2018). Altogether these features will enable to answer questions such as how potential drug candidates affect signal propagation and/or synaptic transmission on a single neuron level (Emmenegger et al., 2019). Finally the high density of electrodes on the CMOS array enables more flexible PDMS microstructure designs with a higher number of single PDMS circuits per array thereby increasing the throughput.

Gaining high reproducibility in experiments using random *in vitro* neural networks has been a big challenge so far (Wagenaar et al., 2006; Napoli et al., 2014; Keren, 2014) and is the reason why the development of central nervous system (CNS)-related drugs has a longer development time and a lower success rate (Gribkoff and Kaczmarek, 2017). Using our directional PDMS microstructures mounted on CMOS arrays enables the design of multiple reproducible neural circuits with predictable activity patterns on a single CMOS array. One main reason why only 5 circuits showed the expected activity pattern is that we did not yet gain full control on the number of neurons per well. We expect that combining our approach with single cell placing techniques (Martinez et al., 2016) to seed ideally only one neuron per well will increase reproducibility and predictability of our small neural networks. Any drug-induced changes on neural network activity will thus be easier to interpret which should enhance the predictive power on how drug candidates will perform in clinical trials (Charvériat et al., 2021).

In this work, we have shown an easy-to-apply method to combine high density CMOS microelectrode arrays with complex PDMS microstructures. Moreover, we developed a Python script to identify the exact location of the electrodes on a CMOS MEA for targeted recording and stimulation. The ability to record from neural networks of any predefined topology on a high resolution CMOS MEA will enable bottom-up neuroscience research to ask new questions on fundamental neuroscience concepts.

## AUTHOR CONTRIBUTIONS

JD and JK conducted the majority of the experiments and the data analysis. SG, TR and SW contributed to the development of imaging methods for the neuron cultures and PDMS microstructures. SJI and CF contributed to the PDMS microstructure design. JH provided SEM images of the CMOS MEA. CF and JV initiated the idea of PDMS microstructure adhesion using an intermediate PDMS gluing step and performed preliminary experiments showing feasibility. AB and CF developed the method and Python scripts to detect the PDMS microstructures on the CMOS MEA electrically. JD and TR wrote the first draft of the manuscript. All co-authors contributed to and approved the manuscript.

## ACKNOWLEDGMENTS

We would like to thank the team from Maxwell Biosystems for their technical support and many helpful discussions.

## SUPPLEMENTAL DATA

**Figure S1** Procedure of imaging the engineered neural networks growing on the CMOS MEA. A coverslip is placed in the culture medium, which is then carefully aspirated, leaving a thin film of culture medium on top of the culture, secured by the coverslip. The MEA is then placed in the headstage, which is flipped and placed in a custom aluminum holder, that can be inserted in an inverted microscope.

**Figure S2** Impedance maps of all 14 CMOS MEAs including information on how many circuits of the microstructures are entirely covered by underlying microelectrodes (“Covered”) and how many circuits are completely open (“Open”), i.e. allow axonal ingrowth.

**Figure S3** Live-dead staining using CMFDA (green, alive) and ethidium homodimer (red, dead) on all 16 CMOS MEAs.

**Movie S1** Video demonstrating the propagation of action potentials in an engineered neural network composed of primary rat neurons at DIV 14.

