## Supplementary figures and images for "Engineered biological neural networks on high density CMOS microelectrode arrays"

### Figure S1

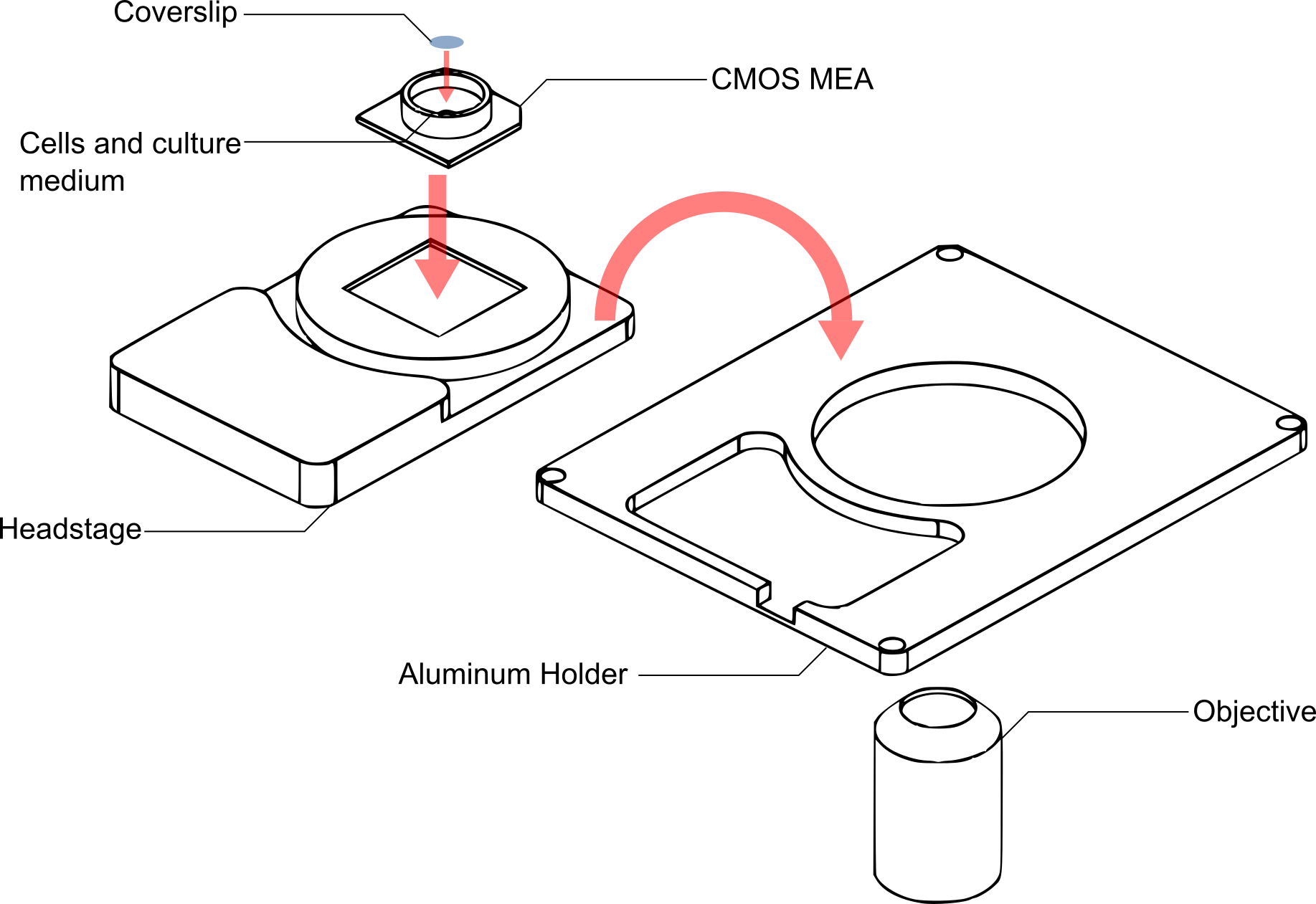

### Figure S2

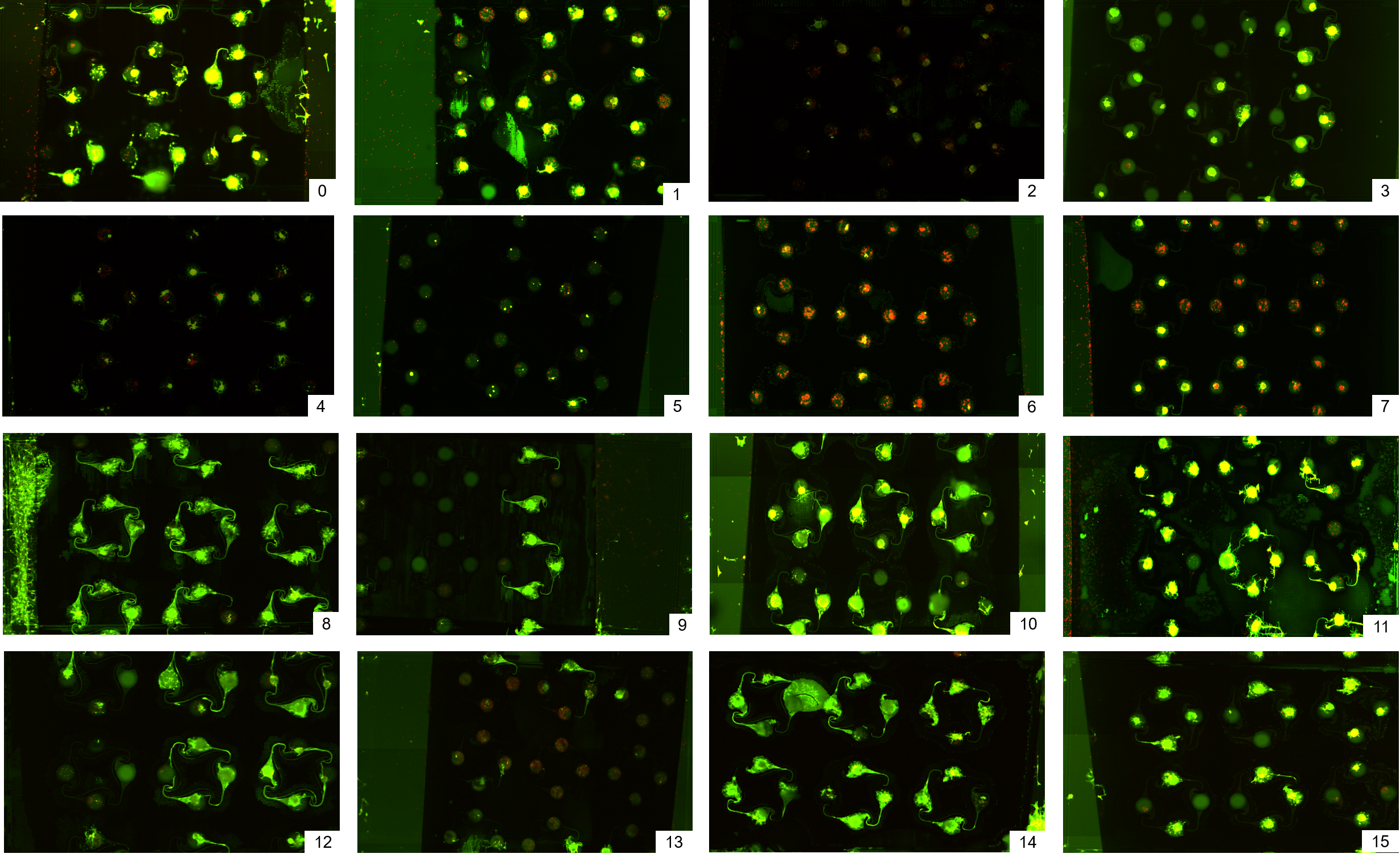

### Figure S3

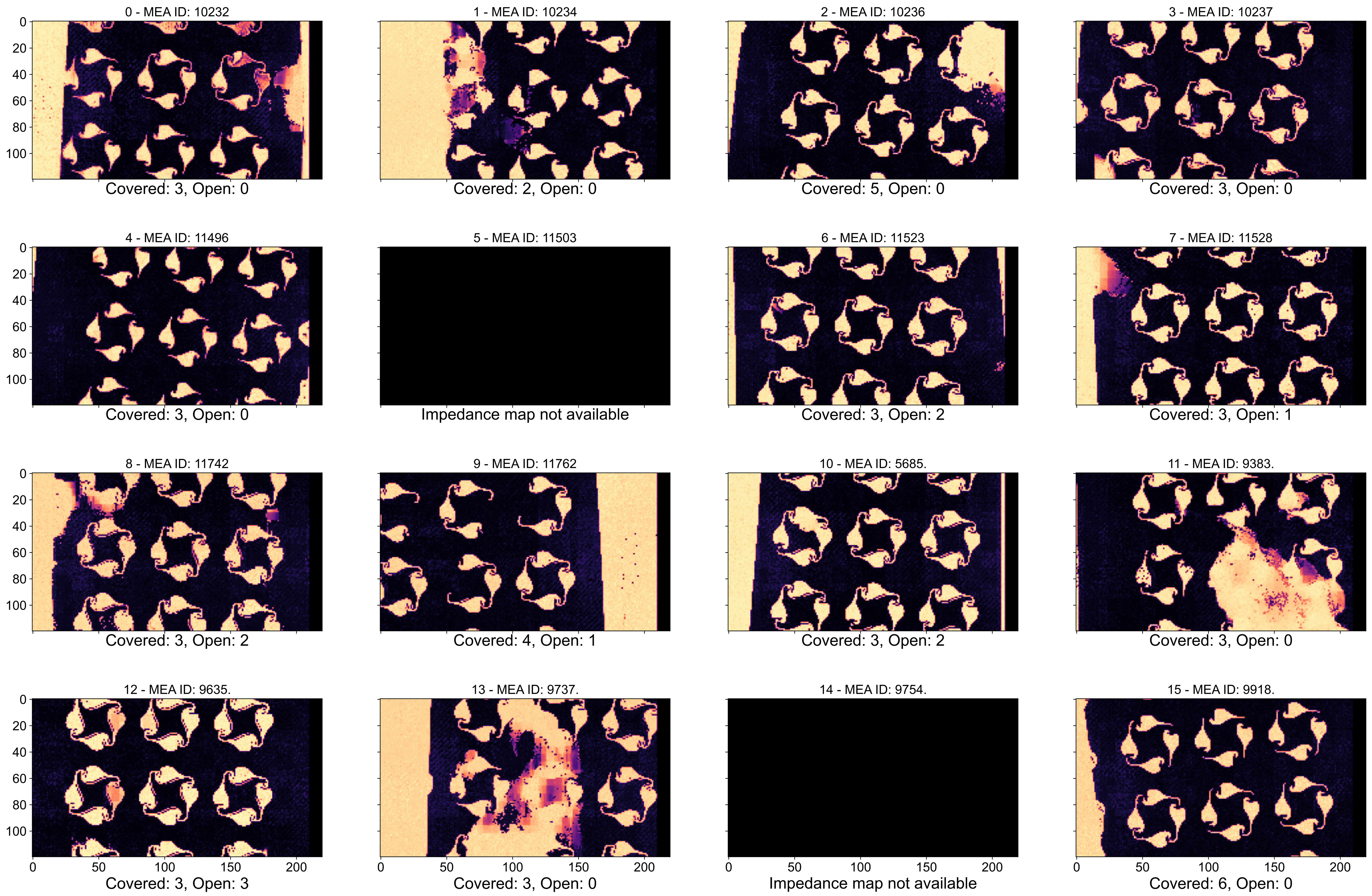
